# PD-1 blockade-driven anti-tumor CD8^+^ T cell immunity requires XCR1^+^ dendritic cells

**DOI:** 10.1101/2020.04.28.066639

**Authors:** Tianyang Mao, Eric Song, Akiko Iwasaki

## Abstract

CD8^+^ T cells are required for effective anti-PD-1 (αPD-1) cancer immunotherapy. Type 1 conventional dendritic cells (cDC1s) bearing XCR1 critically mediate the initiation of protective anti-tumor CD8^+^ T cell responses in mice and humans. However, whether cDC1s contribute to evoking the effector function of CD8^+^ T cells during αPD-1 antibody therapy remains unclear. Here, by deleting cDC1s at the effector phase of αPD-1 therapy, we identify these cells as a crucial innate determinant for effective αPD-1 immunotherapy. αPD-1 treatment unleashed cDC1s to promote anti-tumor CD8^+^ T cell immunity, through the expansion of TCF1^+^ precursors and generation of TIM3^+^ terminally differentiated effectors. Furthermore, tumor cDC1 abundance was predictive of enhanced CD8^+^ T cell infiltration, higher survival, and improved clinical responses to αPD-1 therapy in human cancer patients. Together, this study reveals the requirement for cDC1s in PD-1 blockade therapy, through their ability to elicit CD8^+^ T cell effector responses that mediate tumor control, and highlight cDC1s as an attractive cellular target to be harnessed for novel immunotherapeutics.

## Introduction

Immune checkpoint immunotherapy (ICI) targeting the PD-1/PD-L1 and CTLA-4 pathways has made substantial clinical progress in cancer immunotherapeutics and ushered in the modern era of onco-immunology. Biological factors, such as PD-1/PD-L1 expression^1,2^, tumor neoantigen load^3,4^, deficiency in DNA repair machinery^5,6^, and tumor infiltration by CD8^+^ T cells^7,8^, have proven clinical utility in predicting better responses to αPD-1 antibody therapy. However, the absence of these biomarkers does not preclude clinical efficacy^9,10^. Additionally, although a small fraction of patients derives benefit from durable responses, primary resistance or secondary tumor escape mechanisms rendering immunotherapy ineffective remain an unmet clinical challenge to current ICI^11,12^. Therefore, a better understanding of the molecular and cellular mechanisms that enable and maintain effective clinical responses to cancer immunotherapy is needed.

Tumor-infiltrating lymphocytes (TILs) are subject to local immunosuppression in the tumor microenvironment (TME) critically mediated by the PD-1/PD-L1 pathway^13,14^. Blockade of PD-1/PD-L1 thereby seeks to remove PD-1-mediated inhibition and produce a proliferative response from CD8^+^ TILs. Recent studies suggest that such a proliferative burst originates from a pool of stem-like precursors that express the transcription factor TCF1. TCF1^+^ precursors mediate durable responses to αPD-1 therapy via their self-renewal capacity, and differentiate into TIM3^+^ terminal effectors with greater cytotoxic capacity for further enhancement of tumor restriction^15,16,17,18^. Moreover, an increased ratio of TCF1^+^ to TCF1^−^ CD8^+^ TILs is strongly correlated with clinical responses to ICI and improved survival in human melanoma patients^19^. Thus, the TCF1^+^ precursor/TIM3^+^ effector axis mediates protective anti-tumor immune responses during αPD-1 immunotherapy.

The initiation of anti-tumor CD8^+^ T cell responses requires BATF3-dependent cDC1s, which are a subset of antigen presenting cells (APCs) that specializes in the acquisition, processing, and presentation of tumor antigens^20^. Tumor-infiltrating cDC1s traffic tumor antigens to the tumor draining lymph node (TDLN) in a CCR7-dependent manner where they efficiently cross-prime tumor-specific naive CD8^+^ T cells^21,22,23^. In addition to mediating the induction of endogenous anti-tumor immune responses, recent studies have implicated an important role for cDC1s in the efficacy of checkpoint blockade therapy^24,25,26,27^. In particular, it was shown that tumor-bearing mice that were developmentally deficient in BATF3-dependent DCs failed to respond to αPD-1 or αPD-L1. The lack of an immune response to αPD-1, however, could be due to defective priming of tumor-specific T cells by DCs during early stages of tumor development, and does not reflect the requirement of cDC1s during immunotherapy. Therefore, the role of cDC1s independent of early TDLN priming and their impact on activated, antigen-experienced CD8^+^ T cells during αPD-1 therapy remain unclear. Here we probe the role of cDC1s during the effector stage of αPD-1 cancer immunotherapy.

## Results

### XCR1^+^ cDC1s are required for effective anti-PD-1 immunotherapy against established murine melanoma

We first examined the presence of tumor-infiltrating cDC1s using a melanoma model B16.F10 expressing the lymphocytic choriomeningitis virus glycoprotein epitope GP33-41 (B16.GP33). cDC1s were identified by their surface expression of I-A/I-E, CD11c, XCR1, and CD24, representing a distinct APC compartment in the tumor (Supplementary Fig. 1a,b).

**Figure 1.**
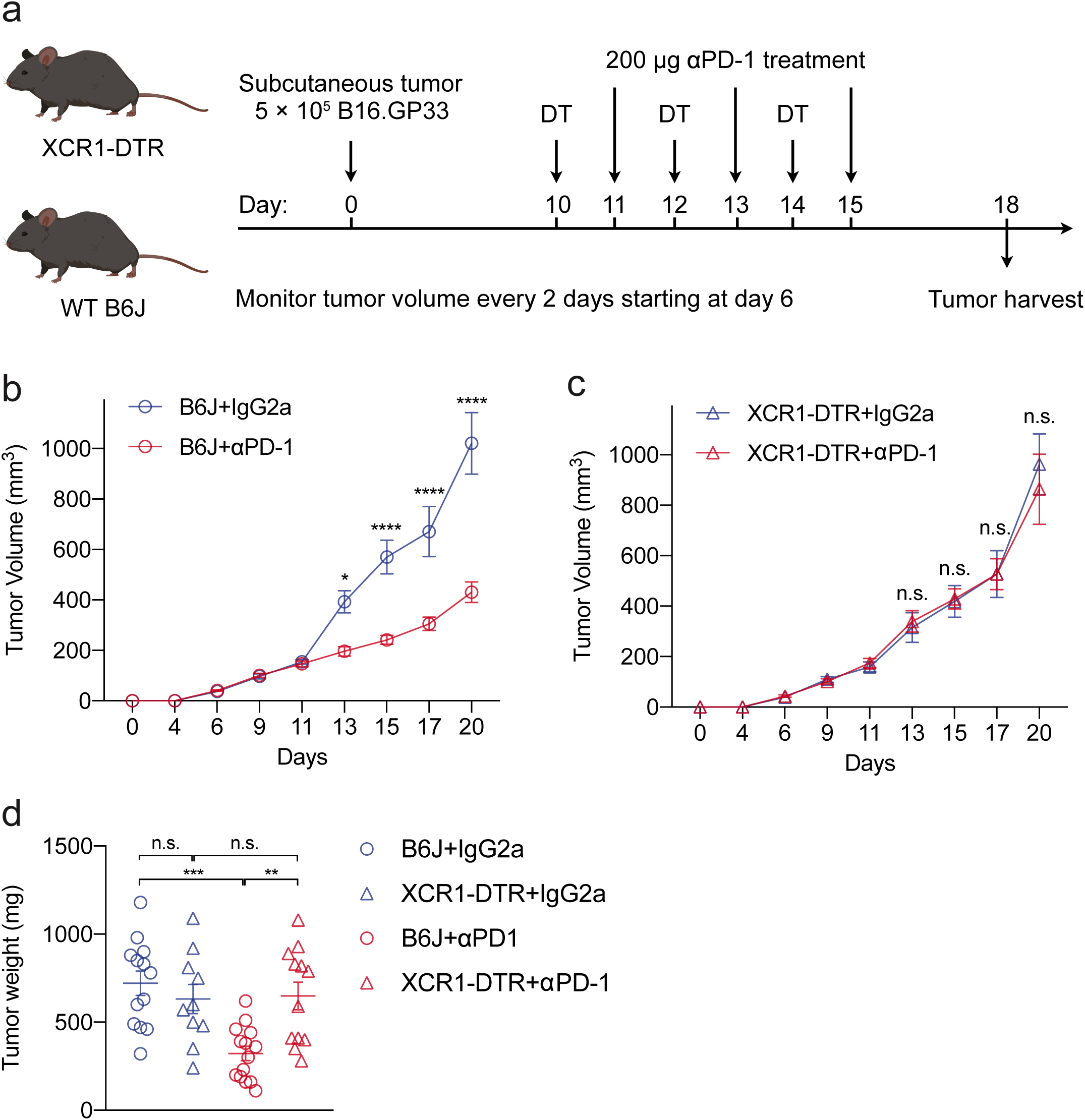
XCR1^+^ cDC1s are required for effective αPD-1 immunotherapy against established murine melanoma. **a**, Treatment scheme: B6J control or XCR1-DTR mice were implanted with 5 × 105 B16.GP33 melanoma. Mice were enrolled in αPD-1 therapy on day 11 post tumor implantation or whenever the tumor volume reached 150 to 200 mm3. 500 ng DT was injected intraperitoneally 24 hours before PD-1 blockade. Both DT and αPD-1 were given every other day for a total of three injections each. **b,c**, Tumor growth curves of B6J (**b**) and XCR1-DTR (**c**) mice treated with either isotype controls or αPD-1 antibodies. **d**, End-point measurement of tumor weight. Data were pooled from three independent experiments with similar results. Mean ± s.e.m., two-way ANOVA (**b,c**) or one-way ANOVA (**d**) followed by Tukey post hoc test; *P ≤ 0.05, **P ≤ 0.01, ***P ≤ 0.001, ****P ≤ 0.0001.

In order to study the in vivo role of cDC1s in αPD-1-mediated CD8^+^ T cell responses while minimizing effects on initial priming, we utilized XCR1-DTR mice in which the human diphtheria toxin receptor (DTR) gene was inserted into the endogenous *Xcr1* locus^28^. Upon diphtheria toxin (DT) injection, tumor-infiltrating and lymph node cDC1s were efficiently depleted in XCR1-DTR mice, but not C57BL/6J (B6J) controls (Supplementary Fig. 2a-f). To assess the requirement of cDC1s in mediating the therapeutic efficacy of PD-1 blockade immunotherapy against melanoma, 5 × 10^5^ B16.GP33 cells were implanted subcutaneously into either XCR1-DTR or B6J mice. Tumor-bearing mice were given three injections of αPD-1 antibodies intraperitoneally every other day starting at day 11 or whenever the tumor size reached 150 to 200 mm^3^. In order to conditionally deplete cDC1s, DT was injected intraperitoneally (i.p.), 24 hours prior to each αPD-1 injection (Fig. 1a). PD-1 blockade significantly delayed tumor growth in B6J mice, whereas untreated mice succumbed to rapid tumor development (Fig. 1b and Supplementary Fig. 3a,c). Notably, when cDC1s were depleted in XCR1-DTR mice, PD-1 blockade was completely rendered ineffective (Fig. 1c and Supplementary Fig. 3b,d). Consistent with tumor growth kinetics, αPD-1 treatment significantly reduced the tumor burden in B6J mice at the experimental end-point compared to untreated controls (Fig. 1d). However, no significant difference in tumor weight was observed between αPD-1- and isotype-treated XCR1-DTR mice (Fig. 1d). Thus, cDC1s are required at the time of αPD-1 antibody therapy.

**Figure 2.**
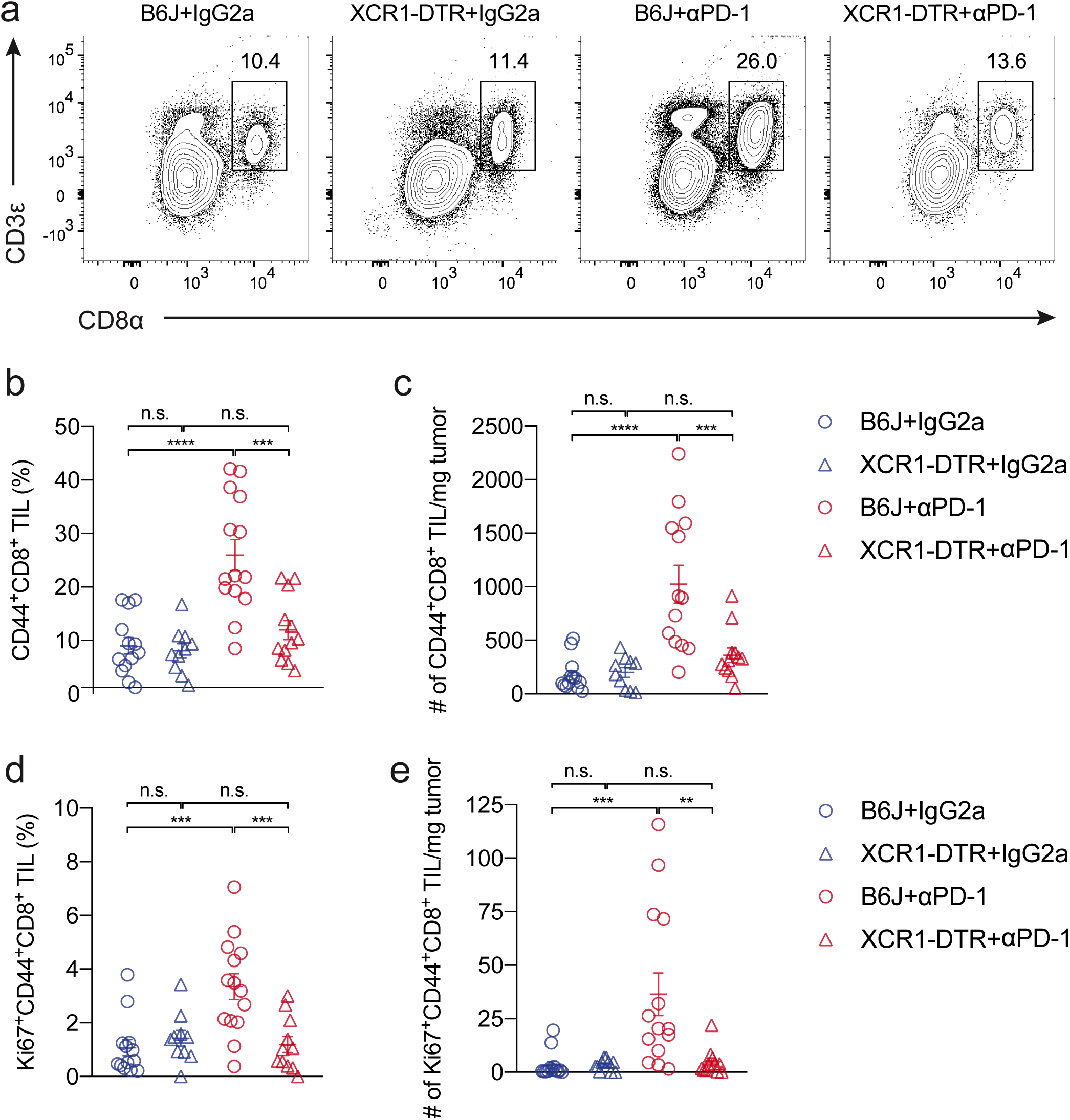
XCR1^+^ cDC1s are required for the robust expansion of intratumoral CD8^+^ T cells after PD-1 blockade. **a-c**, Tumors were dissected and processed for flow cytometric analysis of CD8 T cell responses. Expansion of intratumoral polyclonal CD8^+^ TILs in B16.GP33 in response to αPD-1 treatment was shown as representative flow plot (**a**), frequency of CD45+ immune cells (**b**), and absolute cell number (**c**). **d,e**, Frequency (**d**) and absolute cell number (**e**) of Ki67+ CD44+ CD8^+^ TILs. Data were pooled from three independent experiments with similar results. Mean ± s.e.m., one-way ANOVA (**b,c,d,e**) followed by Tukey post hoc test; *P ≤ 0.05, **P ≤ 0.01, ***P ≤ 0.001, ****P ≤ 0.0001.

**Figure 3.**
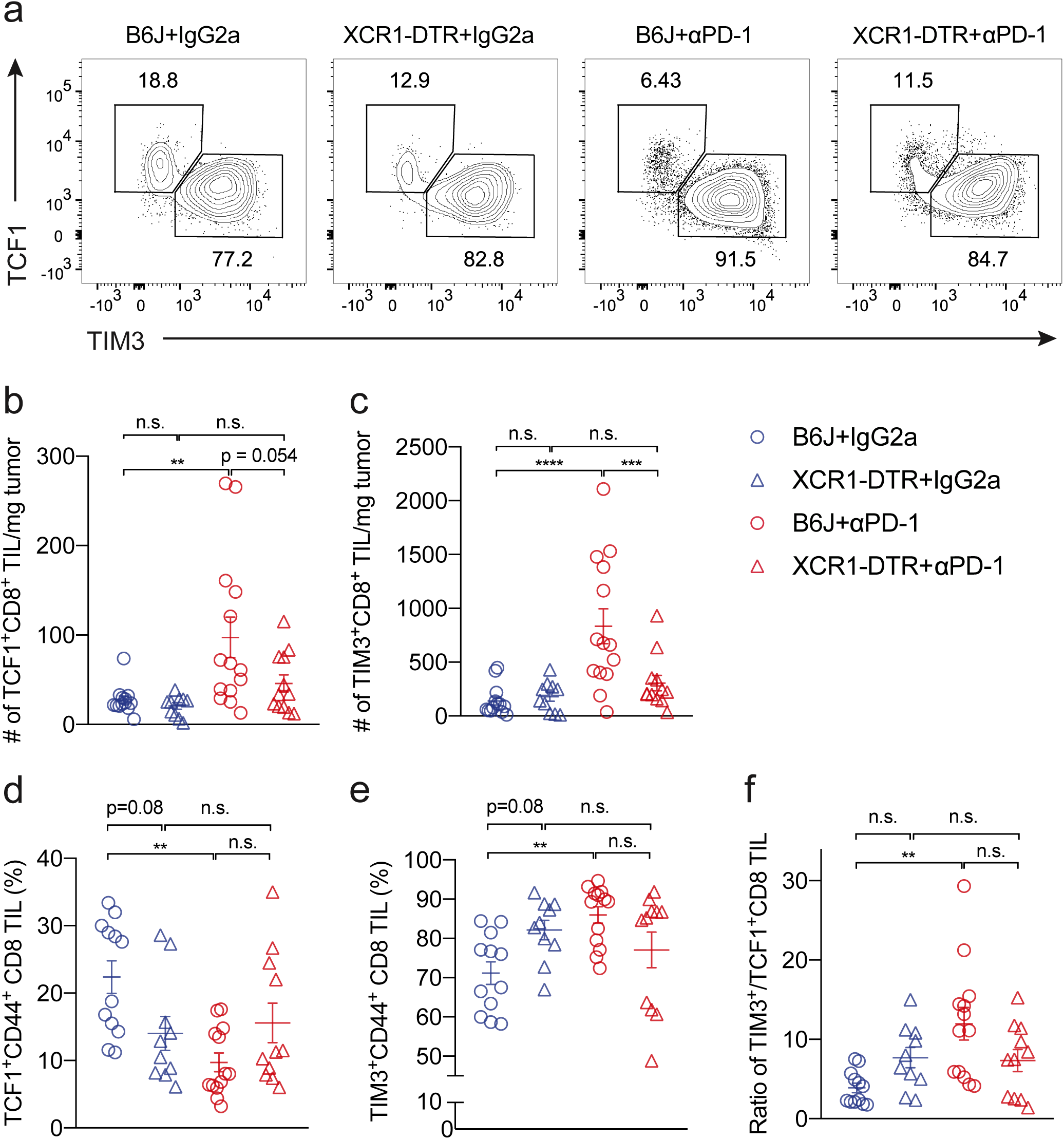
PD-1 blockade-induced terminal differentiation of intratumoral CD8^+^ TILs require XCR1^+^ cDC1s. **a-f**, Representative flow plot (**a**) and summary of absolute number and relative frequency of polyclonal TCF1^+^ (**b,d**) and TIM3^+^ (**c,e**) CD8^+^ TILs in B16.GP33 tumor. Differentiation of TCF1^+^ progenitors into TIM3^+^ terminal effectors was also shown as ratio of TCF1^+^/TIM3^+^ CD8^+^ TILs (**f**). Data were pooled from three independent experiments with similar results. Mean ± s.e.m., one-way ANOVA (**b,c,d,e,f**) followed by Tukey post hoc test; *P ≤ 0.05, **P ≤ 0.01, ***P ≤ 0.001, ****P ≤ 0.0001.

### XCR1^+^ cDC1s are required for the robust expansion of intratumoral CD8^+^ T cells after PD-1 blockade

Next, we examined whether cDC1s are required to elicit CD8 T cell responses following PD-1 blockade. Consistent with previous clinical observations and mouse studies, PD-1 blockade significantly induced the accumulation of CD8^+^ TILs within the tumor in B6J mice (Fig. 2a-c and Supplementary Fig. 4). These cells were defined by the surface expression of CD44, indicative of their antigen-experienced state. However, increases in CD8^+^ TILs were completely abolished in the absence of cDC1s (Fig. 2a-c). Intratumoral accumulation of CD8^+^ TILs can be partially explained by enhanced proliferation, as PD-1 blockade significantly increased both frequency and absolute number of Ki67^+^ CD8^+^ TILs (Fig. 2d-e). Thus, cDC1s are required to induce the activation and proliferation of CD8^+^ TILs in response to αPD-1.

**Figure 4.**
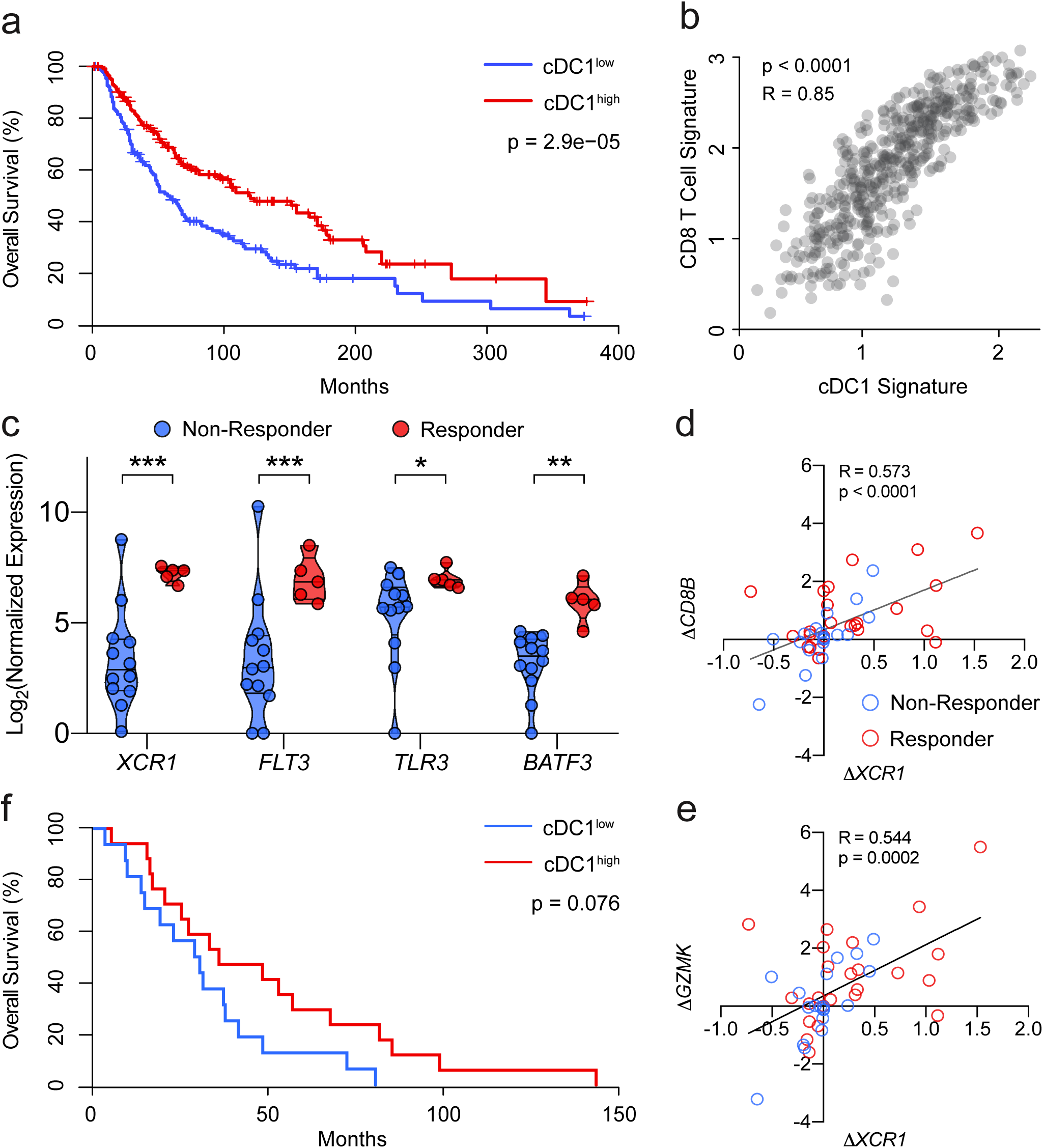
cDC1 signature is associated with overall survival, clinical efficacy, and CD8^+^ T cell responses to αPD-1 immunotherapy in human cancer patients. **a**, Association between cDC1 signature expression and overall survival in TCGA skin cutaneous melanoma (SKCM) patients. **b**, Correlation between expression of cDC1 signature and effector CD8 T cell signature in TCGA SKCM cohorts. cDC1 signature is defined as *XCR1*, *BATF3*, *WDFY4*, *FLT3*, *IRF8*, *TLR3*, and *CLEC9A*. Effector CD8 T cell signature is defined as *CD8A*, *CD8B*, *GZMA*, *GZMB*, *GZMK*, and *PRF1*. **c**, Association between cDC1 signature expression and clinical responses to αPD-1 therapy from metastatic melanoma patient biopsies (median-time 1.4 months after αPD-1 treatment). **d,e**, Correlation between changes in the expression of XCR1 and that of CD8 T cell markers *CD8B* (**d**), or *GZMK* (**e**) after αPD-1 treatment in melanoma patients. **f**, Association between pre-treatment tumor cDC1 signature expression and survival in clear cell renal cell carcinoma patients treated with αPD-1 antibody therapy. Mean ± s.e.m., log-rank Mantel–Cox test (**a,f**), two tailed Pearson’s correlation (**b,d,e**), two-way ANOVA followed by FDR correction (**c**); *P ≤ 0.05, **P ≤ 0.01, ***P ≤ 0.001, ****P ≤ 0.0001.

### PD-1 blockade-induced terminal differentiation of intratumoral CD8^+^ TIL require XCR1^+^ cDC1s

TCF1^+^ CD8^+^ TILs serve as a reservoir to sustain CD8^+^ T cell responses, and are preferentially targeted by PD-1 blockade for expansion and generation of TIM3^+^ effectors16. Since durable responses to αPD-1 therapy require a TCF1^+^ stem-like CD8^+^ T cell population, we sought to probe the effect of cDC1s on these precursor cells and their terminally differentiated progeny. We observed a significant increase in the absolute number of both TCF1^+^ and TIM3^+^ CD8^+^ TILs after αPD-1 treatment (Fig. 3a-c). Additionally, PD-1 blockade shifted the distribution of CD8^+^ TILs from TCF1^+^ cells to those with a TIM3^+^ phenotype, suggesting enhanced differentiation of stem-like precursors into terminal effectors. In the absence of cDC1s, however, αPD-1 was no longer capable of expanding the pool of precursors or terminal effectors (Fig. 3a-c). The ability for TCF1^+^ precursors to form TIM3^+^ terminal effectors was also completely abrogated in the absence of cDC1s (Fig. 3d-f). Concordant with superior effector functions of TIM3^+^ CD8^+^ TILs compared to the precursors, αPD-1 induced the production of IFNγ and TNFα by CD8^+^ TILs, which was also dependent on the presence of cDC1s (Supplementary Fig. 5a,b). Together, these data indicated that cDC1s are required for the expansion of precursor CD8^+^ TILs and their differentiation into terminal effectors capable of secreting IFNγ and TNFα in αPD-1 antibody therapy.

### cDC1 signature is associated with overall survival, clinical efficacy, and CD8^+^ T cell responses to anti-PD-1 therapy in human cancer patients

Finally, we examined the association of cDC1 tumor presence in cancer patients with survival and CD8 T cell infiltration. We first defined a cDC1 signature, comprising *XCR1*, *FLT3*, *TLR3*, *BATF3*, *CLEC9A*, *WDFY4*, and *IRF8*, to assess the extent of tumor infiltration by cDC1s^9^. Through the analysis of The Cancer Genome Atlas (TCGA) database, we found that the level of cDC1 signature expression positively correlated with overall survival in skin cutaneous melanoma (SKCM) patients not treated with ICI^29^ (Fig. 4a and Supplementary Fig. 6a-f). Superior survival in cDC1^high^ patients can be partially explained by stronger effector CD8^+^ T cell responses within the tumor, as the expression of the cDC1 signature highly correlated with that of signature genes defining effector CD8^+^ T cells, including *CD8A*, *CD8B*, *GZMA*, *GZMB*, *GZMK*, and *PRF1* (Fig. 4b and Supplementary Fig. 7a-f). These results suggested that cDC1s drive endogenous anti-tumor CD8^+^ T cell responses in SKCM patients.

We next extended our analyses of cDC1 and CD8^+^ T cell responses to cancer immunotherapy. In metastatic melanoma patients, the expression level of various cDC1 genes from clinical biopsies (obtained median-time 1.6 months after the treatment) were positively correlated with responsiveness to αPD-1 immunotherapy^30^ (Fig. 4c). Furthermore, using data from a different melanoma cohort, we found that an increase in *XCR1* expression was positively correlated with that of effector CD8^+^ T cell markers, regardless of the therapeutic efficacy^12^ (Fig. 4d,e and Supplementary Fig. 8a-f). However, bigger changes in the expression of both *XCR1* and CD8^+^ T cell marker genes were observed in responders. Lastly, pre-treatment expression level of cDC1 signature genes from clear cell renal cell carcinoma and metastatic patients was predictive of clinical responses to αPD-1 therapy^6,31^ (Fig. 4f and Supplementary Fig. 9a,b). Therefore, intratumoral cDC1s levels correlate strongly with patient survival in general, and with anti-PD1 antibody therapy success. Furthermore, intratumoral cDC1 signature appears to serve as a biomarker for predicting successful cancer immunotherapy.

## Discussion

In this study, we examined the role of cDC1s in the effector function of CD8^+^ T cells that enable effective αPD-1 immunotherapy. Temporal ablation of cDC1s abrogated αPD-1- mediated tumor growth inhibition. Mechanistically, cDC1s promoted the proliferative response of intratumoral CD8^+^ TILs, expanded the pool of TCF1^+^ stem-like precursors, and induced the generation of TIM3^+^ terminal effectors during αPD-1 therapy. Furthermore, the expression of cDC1 signature genes positively correlated with overall survival, clinical efficacy, and CD8^+^ T cell responses to αPD-1 therapy in human cancer patients. Collectively, our data identified the crucial role for cDC1s in evoking CD8^+^ T cell responses during αPD-1 therapy and highlighted the importance of therapeutic targeting of cDC1s for optimal induction of anti-tumor CD8^+^ T cell immunity.

These data suggest that the ability of cDC1s to induce intratumoral CD8^+^ T cell responses is normally ineffective and relies on inhibition of PD-1/PD-L1 to unleash their stimulatory functions. Recent work has demonstrated a critical role for the CD28/B7 pathway in the proliferative rescue of CD8^+^ T cells and the enhanced control of tumor growth by PD-L1 blockade in mice^32^. Moreover, it was shown that intratumoral cDC1-derived IL-12 activates CD8^+^ TILs and contributes for effective PD-1 blockade therapy^27^. Given the various avenues through which cDC1-CD8 T cell crosstalk could be established, future studies will need to determine precisely how cDC1s elicit robust CD8 T cell responses during αPD-1 therapy.

The concept of intratumoral immune activation by PD-1 blockade has greatly advanced our understanding of the immunological basis for successful immunotherapy. However, observations of enhanced T cell infiltration and function in tumors after αPD-1 therapy do not formally prove the importance of intratumoral TIL reinvigoration in ICI therapy. Consistent with this idea, recent scRNA-seq studies in cancer patients have shown that the majority of tumor-specific TILs that expanded in response to PD-1 blockade was not initially detected in the tumor but rather had a peripheral origin and can be detected in the blood^33,34^. Although PD-1/PD-L1 engagement at the TIL-tumor interface still remains a critical barrier to overcome for the elicitation of protective CD8^+^ T cell responses, these studies raised the possibility that tumor-specific CD8^+^ T cells can respond peripherally, presumably through the help of PD-L1-expressing cells, and can be readily recruited into the tumor during αPD-1 treatment. As cDC1s can be found both in the tumor and TDLN, future studies are needed to pinpoint the tissue compartment(s) in which cDC1-mediated immune activation occur in response to αPD-1.

Innate control of adaptive immune responses is a well-established paradigm^35^. Through the study of immunological mechanisms behind effective PD-1 blockade, here we provide data to extend this paradigm to cancer immunotherapy. Furthermore, our study provides explanations as to why therapeutic modulations that target DCs for expansion and activation, as well as their synergy with ICI, demonstrated clinical efficacy. Finally, our results lay the foundation for novel strategies that aim to improve the ability of cDC1s for optimal anti-tumor CD8^+^ T cell responses.

## Materials and Methods

### Mice

C57BL/6J mice were bred and maintained in our specific-pathogen free animal colony. XCR1-DTR mice were obtained from T. Kaisho (Wakayama University). All procedures used in this study complied with federal guidelines and the institutional policies of the Yale School of Medicine Animal Care and Use Committee.

### Cell lines

B16.GP33 cells were a gift from S. Kaech (Salk Institute). Cells were cultured in complete DMEM (4.5 g/L D-Glucose, L-Glutamine, 1% penicillin/streptomycin, Gibco) supplemented with 10% FBS (Sigma) and regularly tested for mycoplasma using the MycoAlertTM Mycoplasma Detection Kit (LONZA).

### Treatments

DT (List Biological Laboratories) was reconstituted as 1 mg/mL stock solution in PBS (Sigma). 25 ng/g body weight DT was injected intraperitoneally into the mice on day 10, 12, and 14. 200 µg αPD-1 (clone 29 F.1A12, BioXCell) or 200 µg isotype-matched control antibody (Rat IgG2a, BioXCell) were given intraperitoneally on day 11, 13, and 15. DT treatment regimen was designed based on previous studies showing that cDC1s were efficiently depleted on day one and day two after DT injection, and began to recover at day four ^28^.

### Tumor implantation, measurement, and processing

Mice were subcutaneously implanted with 5 × 10^5^ B16.GP33 cells in sterile PBS. Tumor volume was measured as (Width^2^ × Length)/2 starting at day 6 every 2 to 3 days using a digital caliper. At the experimental end point (day 18 to 20), tumors were dissected, mechanically minced and digested with collagenase D (1 mg/mL, Sigma) and DNase I (60 µg/mL, Sigma) in complete RPMI-1640 media (Gibco) for 30 min at 37 °C. Tumor samples were filtered to single cell suspension and further treated with ACK buffer to remove red blood cells. Tumor draining lymph nodes (inguinal and axillary) were collected at the same time, mechanically minced and filtered to single-cell suspension. Tumor and draining lymph node samples were then counted using an automated cell counter (Thermo Fisher Scientific).

### In vitro T cell stimulation

In vitro T cell simulation assays were performed using complete RPMI-10 supplemented with 10% FBS, 1% penicillin/streptomycin (Gibco), 1 mM sodium pyruvate (Gibco), 100 µM non-essential amino acids (Gibco), 20 mM HEPES (Gibco), and 50 uM β-mercaptoethanol (Sigma). Cells were stimulated with 200 µL 1× eBioscience Cell Stimulation Cocktail (ThermoFisher) without protein transporter inhibitor for 1 hr at 37 °C. An additional 50 µL of 5× eBioscience Cell Stimulation Cocktail with protein transporter inhibitor (ThermoFisher) was added and cells were treated for an additional 4 hr at 37 °C.

### Flow Cytometry

Tumor and draining lymph node samples were blocked with anti-mouse CD16/32 antibodies (BioXCell) for 20 min at 4 °C and stained with surface antibodies (Supplementary Table 1) for 30 to 60 min at 4 °C. Dead cells were excluded with Fixable Aqua added together with anti-CD16/32 (ThermoFisher). For intracellular staining, cells were fixed with 2% formaldehyde (J.T. Baker) for 45 min at 4 °C and then permeabilized with either 1 × Permeabilization Buffer (transcriptional factor staining, ThermoFisher) or 1 × Perm/Wash Buffer (cytokine staining, BD Biosciences) for 10 min at room temperature. Permeabilized cells were stained with intracellular antibodies (Supplementary Table 1) for 30 min at 4 °C. Data were collected with BD LSR II flow cytometer (BD Bioscience). Analysis was performed using FlowJo software (v10, TreeStar).

Cell staining was performed using the following fluorophore-conjugated antibodies: CD45.2 (104), CD19 (6D5), NK1.1 (PK136), TCRβ (H57-597), CD3e (145-2C11), CD8α (53-6.7), CD44 (IM7), Ki67 (16A8), IFNγ (XMG1.2), TNFα (MP6-XT22), TIM3 (RMT3-23), CD11c (N418), I-A/I-E (M5/114.15.2), XCR1 (ZET), CD24 (M1/69), CD11b (M1/70), CD64 (X54-5/7.1), Ly6C (HK1.4), and TCF1 (C63D9).

### Human data analyses

For RNA-seq data, gene expression was normalized to transcripts-per-million (TPM) or fragments per kilobase per million mapped reads (FPKM). For NanoString data, raw expression values were background corrected and RNA-content normalized using probe sets targeting housekeeping genes (*ABCF1*, *GUSB*, *TBP*, and *TUBB*). Log2-transformed data were used for downstream analyses. All TCGA analyses were performed using the web server GEPIA2. Survival analysis of the TCGA SKCM patients was performed based on the level of cDC1 signature expression, defined as the average expression of *XCR1*, *FLT3*, *TLR3*, *BATF3*, *CLEC9A*, *WDFY4*, and *IRF8*, and the patients were dichotomized based on median value of cDC1 signature expression. Pair-wise gene correlation analysis was performed between cDC1 signature and effector CD8 T cell signature, which was defined as the average expression of *CD8A*, *CD8B*, *GZMA*, *GZMB*, *GZMK*, and *PRF1.*

Log2-transformed fold change (Δ) values were calculated for correlation analysis between changes in expression of *XCR1* and that of CD8 T cell markers before and after PD-1 blockade in melanoma patients. For survival analysis, clear cell renal cell carcinoma and metastatic melanoma patients receiving αPD-1 therapy were dichotomized based on median level of cDC1 signature expression to assess the association between tumor cDC1 abundance prior to ICI and clinical responses.

### Statistical analyses

Statistical analyses were conducted in Prism 8 (GraphPad Software). All results were expressed as mean ± s.e.m. Means between two groups were compared with unpaired two-tailed Student’s t tests. Means among more than two groups were compared with one-way or two-way ANOVA followed by Tukey post hoc test. Survival curves were compared using two-sided log-rank Mantel-Cox test or Gehan-Breslow-Wilcoxon test. Correlation analysis was performed using two-tailed Pearson’s correlation.

## Acknowledgements

The authors gratefully acknowledge members of the Iwasaki lab for helpful feedback, Melissa Linehan for logistical and technical assistance, Huiping Dong for mouse breeding, and Orr-El Weizman for critical reading of the manuscript. This study was supported by National Institutes of Health grants T32AI007019 (YIITP training grant), T32GM007205 (MSTP training grant), F30CA239444 (to E.S.), AI054359 (to A.I.), and AI127429 (to A.I.). A.I. is an investigator of the Howard Hughes Medical Institute. The results are in part based upon data generated by the TCGA Research Network (https://www.cancer.gov/tcga).

## Author Contributions

A.I. supervised the research; T.M. and A.I. designed the experiments, analyzed data, and prepared the manuscript; T.M. performed experiments; T.M. and E.S. analyzed published human datasets.

---

**Supplementary Figure 1.**
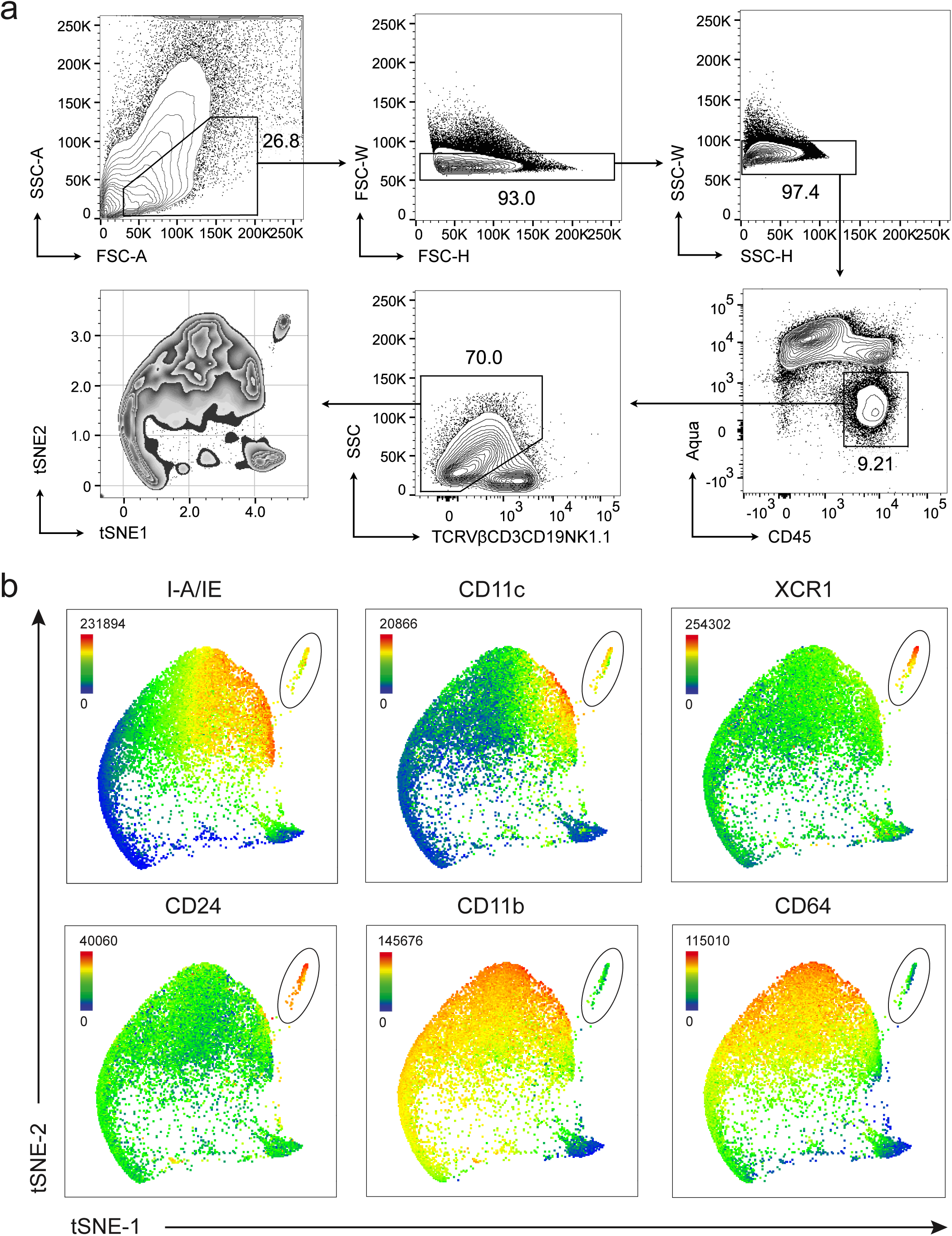
XCR1^+^ cDC1s constitute a unique APC compartment in murine melanoma. **a**, Gating schema for flow cytometric analysis of intratumoral myeloid lineages. **b**, Expression of various surface markers on tumor-infiltrating cDC1s (circled) in B16.GP33. Data were projected into tSNE space. Immune cells expressing CD3ε, TCRVβ, CD19, or NK1.1 were excluded from the analysis.

**Supplementary Figure 2.**
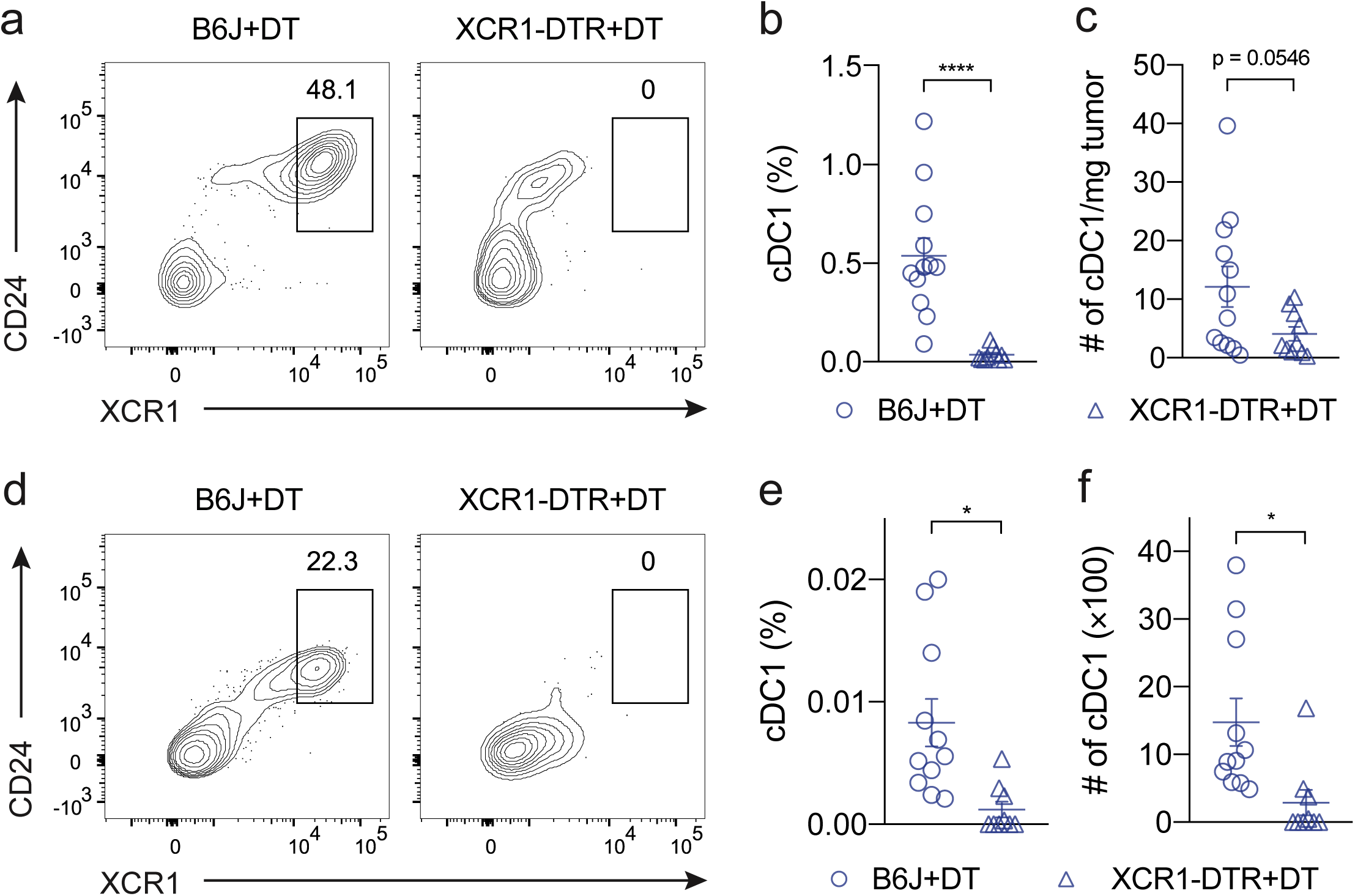
XCR1^+^ cDC1s can be targeted for efficient depletion. **a-f**, cDC1s in tumor (**a-c**) and TDLN (**d-f**) in XCR1-DTR mice were depleted through intraperitoneal injection of 25 ng/g body weight DT three times every other day. cDC1s were defined as CD45+Lineage-CD64-Ly6C-CD11b-MHCII+CD11c+XCR1^+^CD24+. Depletion efficiency was shown as representative flow plot (**a,d**), percentage of CD45+ immune cells (**b,e**), and absolute cell number (**c,f**). Data were pooled from three independent experiments with similar results. Mean ± s.e.m., two-sided Student’s t-test (**b,c,e,f**); *P ≤ 0.05, **P ≤ 0.01, ***P ≤ 0.001, ****P ≤ 0.0001.

**Supplementary Figure 3.**
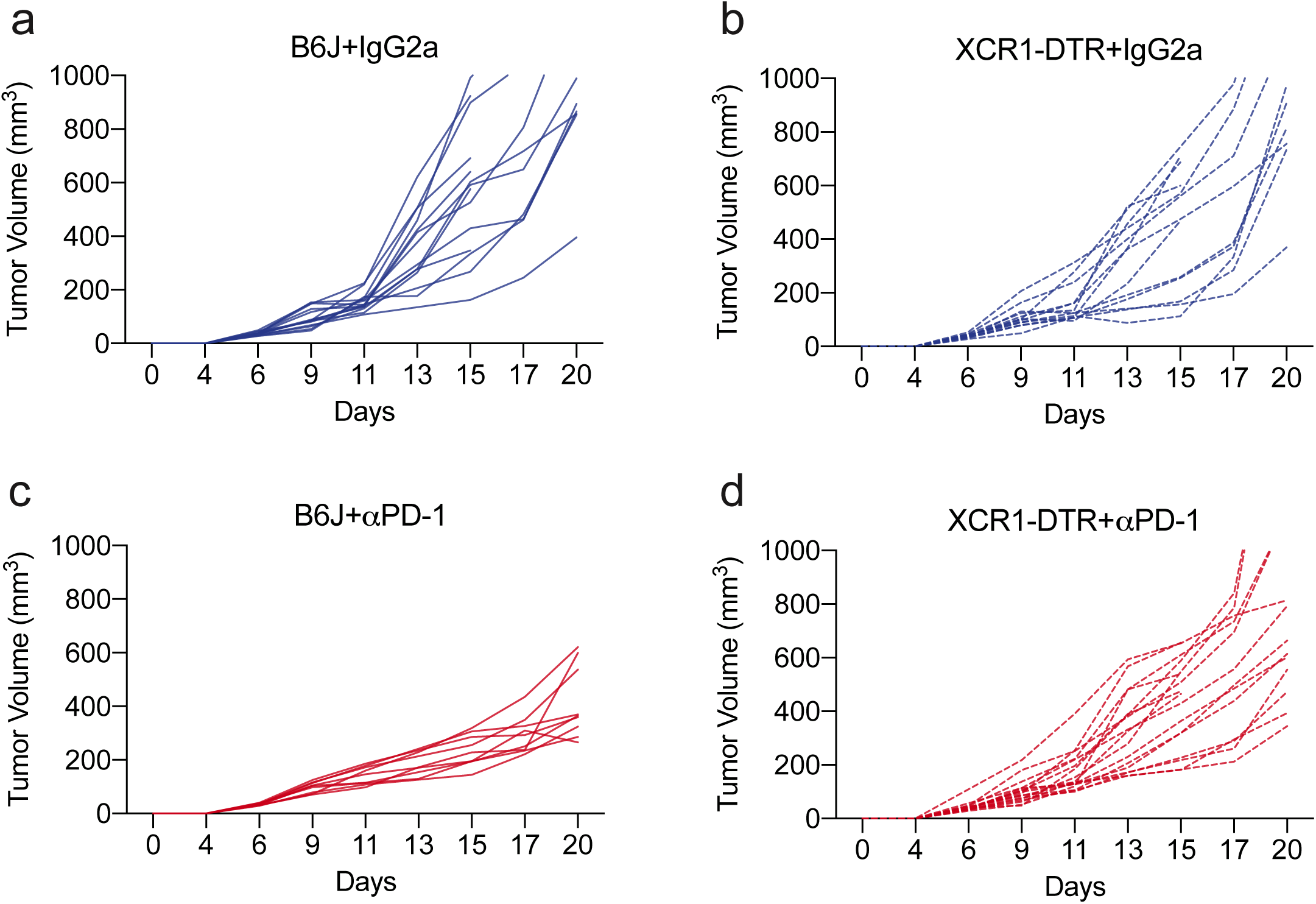
Tumor growth curves of individual mice from Figure 1. Tumor growth curves of B6J+IgG2a (**a**), XCR1-DTR+IgG2a (**b**), B6J+αPD-1 (**c**), and XCR1-DTR+αPD-1 (**d**) group. Tumor volume was measured as (Width2 × Length)/2 starting on day 6 every 2 to 3 days using a digital caliper.

**Supplementary Figure 4.**
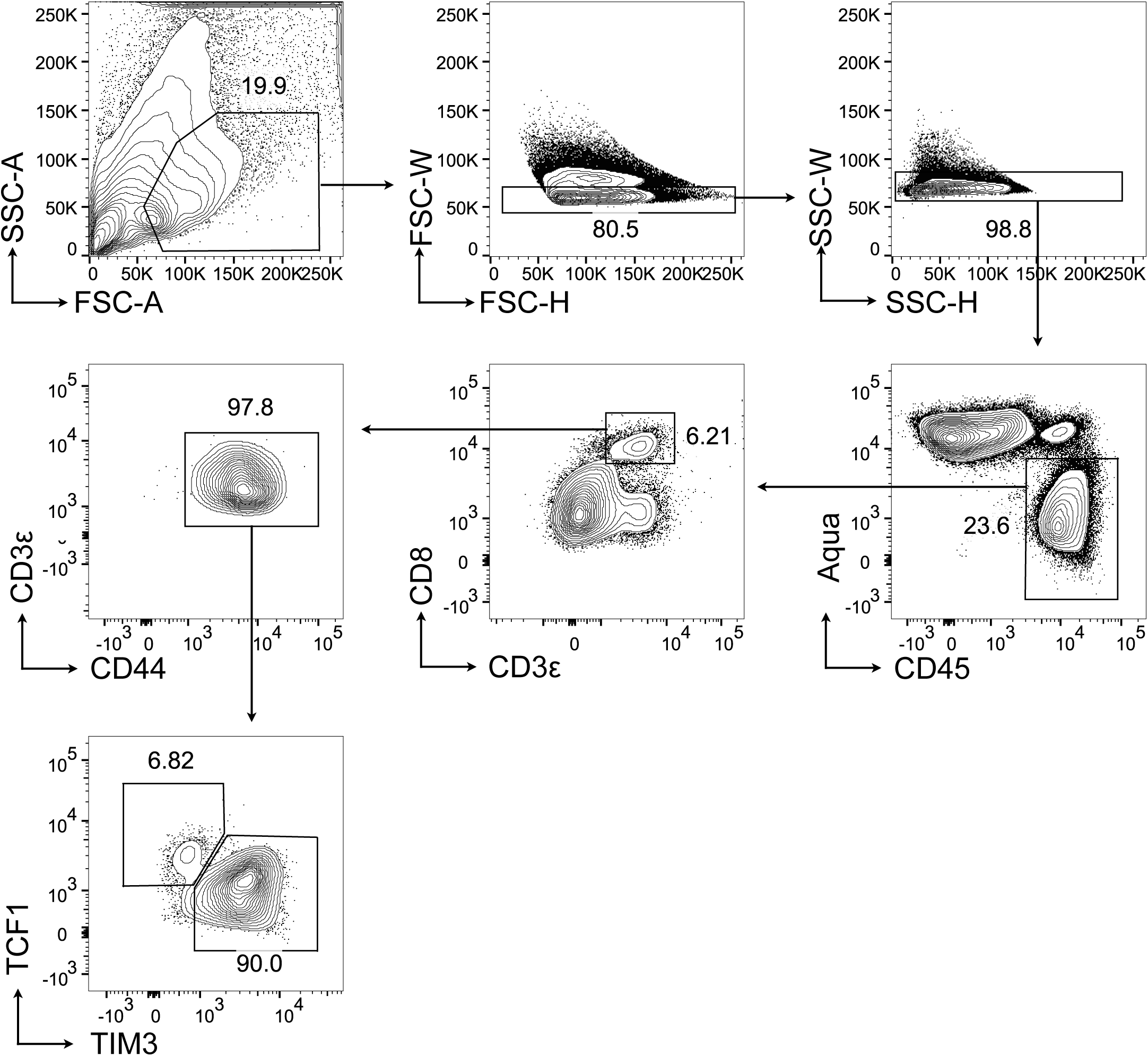
Gating schema for flow cytometric analysis of intratumoral T cells.

**Supplementary Figure 5.**
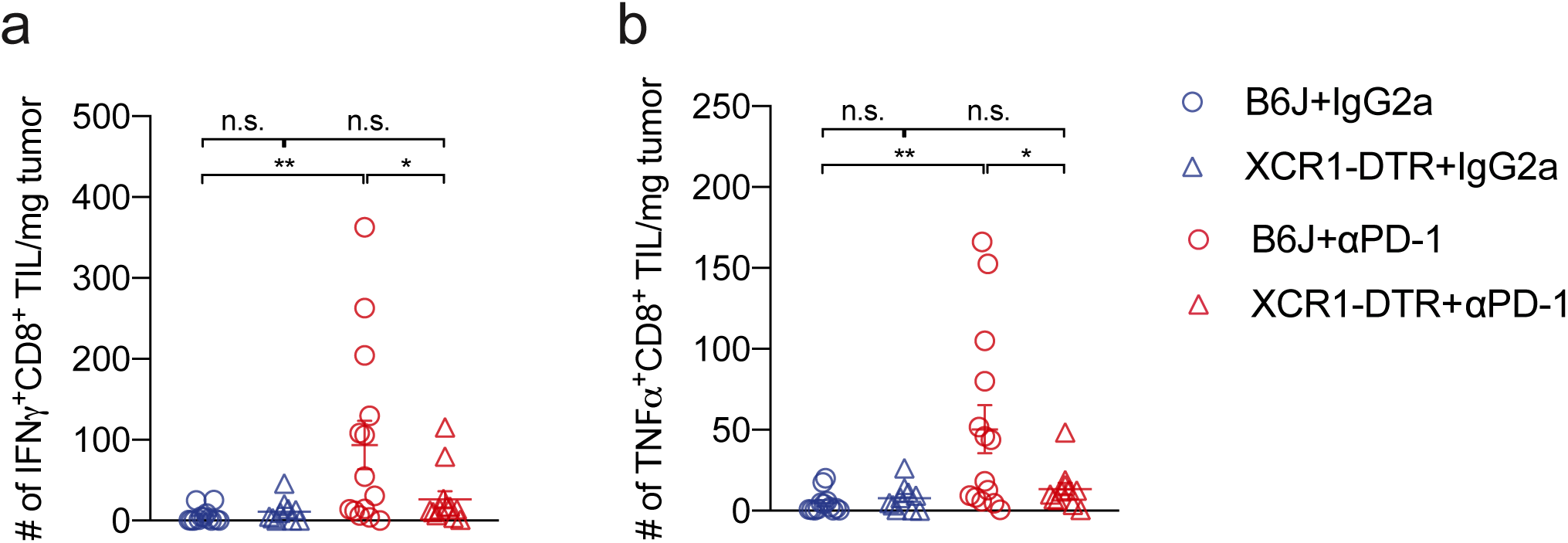
PD-1 blockade-induced functional reinvigoration of intratumoral CD8^+^ T cells require XCR1^+^ cDC1s. **a,b**, Absolute cell number of IFNγ+ (**a**) and TNFα+ (**b**) CD8^+^ TILs. Data were pooled from three independent experiments with similar results. Mean ± s.e.m., one-way ANOVA (**a,b**) followed by Tukey post hoc test; *P ≤ 0.05, **P ≤ 0.01, ***P ≤ 0.001, ****P ≤ 0.0001.

**Supplementary Figure 6.**
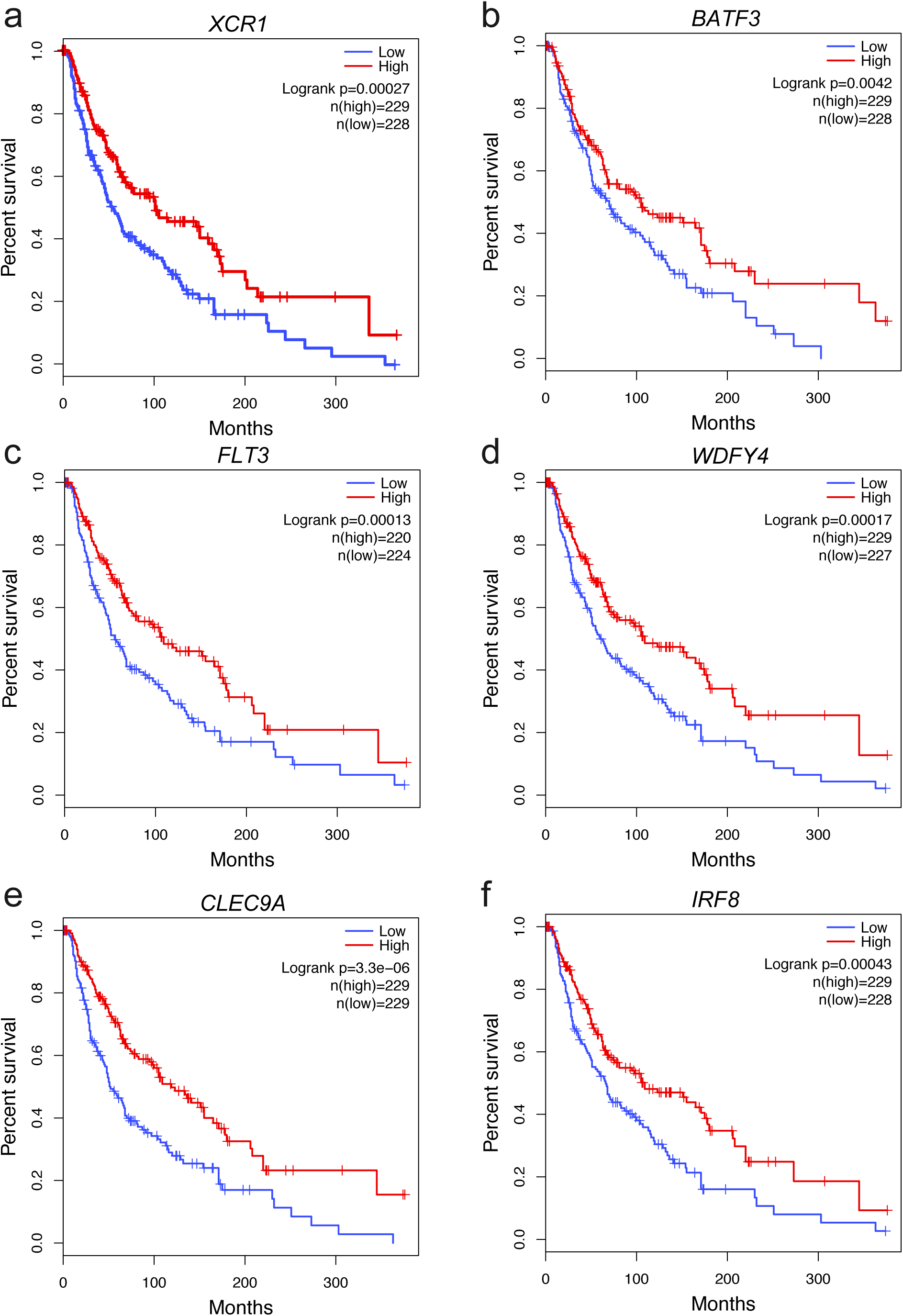
Survival analysis of cDC1 signature gene expression in TCGA SKCM patients. Association between expression of cDC1 markers gene *XCR1* (**a**), *BATF3* (**b**), *FLT3* (**c**), *WDFY4* (**d**), *CLEC9A* (**e**), and *IRF8* (**f**), and overall survival of TCGA SKCM patients. Patients were dichotomized based on median log2-transformed TPM values of each gene. High and low expression groups were compared using Log-rank Mantel–Cox test (**a-f**). Number of patients used for the analysis was indicated in each survival plot.

**Supplementary Figure 7.**
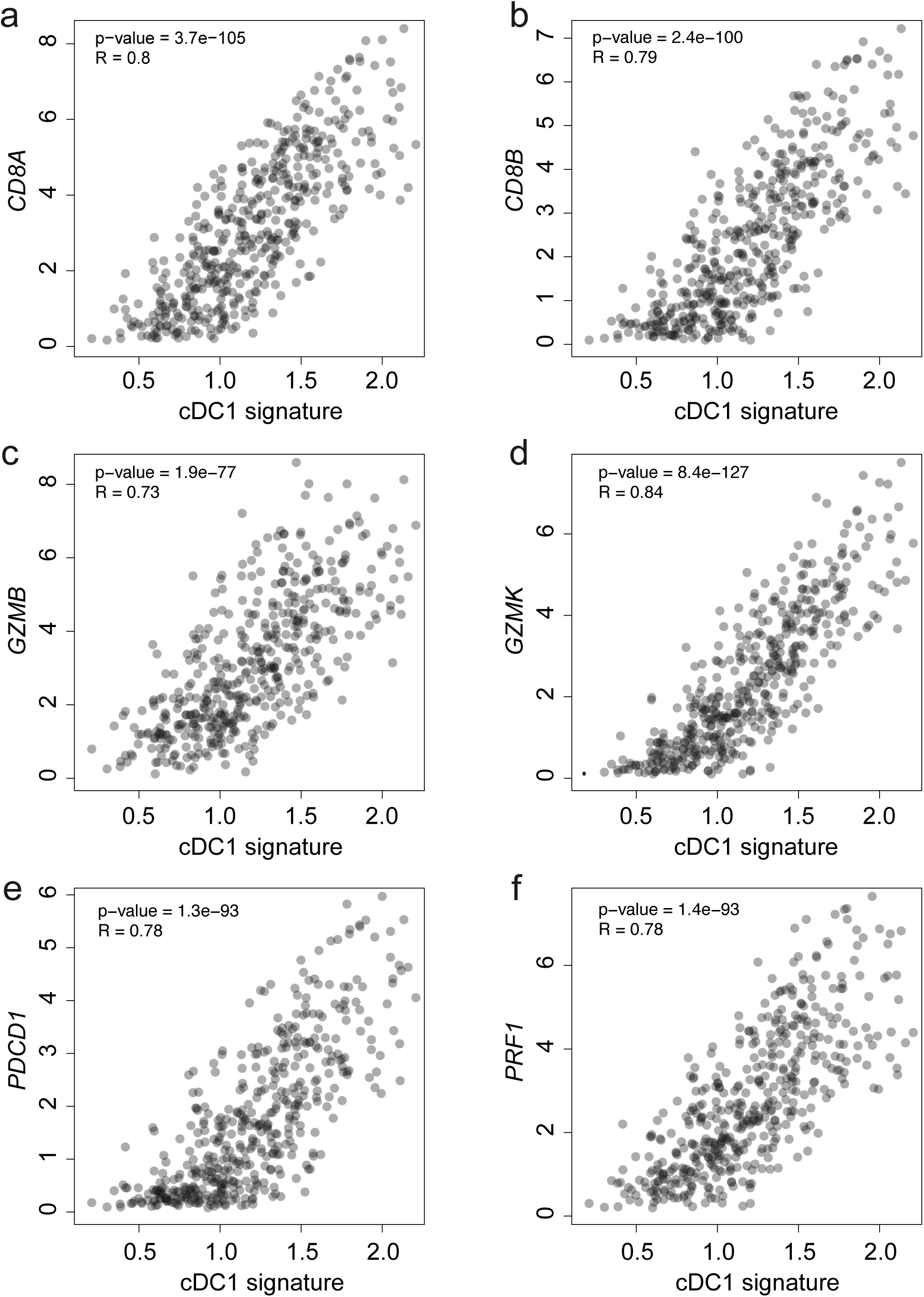
Correlation analysis between cDC1 signature expression and effector CD8 T cell markers in TCGA SKCM patients. Pair-wise Pearson’s correlation was calculated between cDC1 signature and effector CD8 T cell markers *CD8A* (**a**), *CD8B* (**b**), *GZMB* (**c**), *GZMK* (**d**), *PDCD1* (**e**), and *PRF1* (**f**).

**Supplementary Figure 8.**
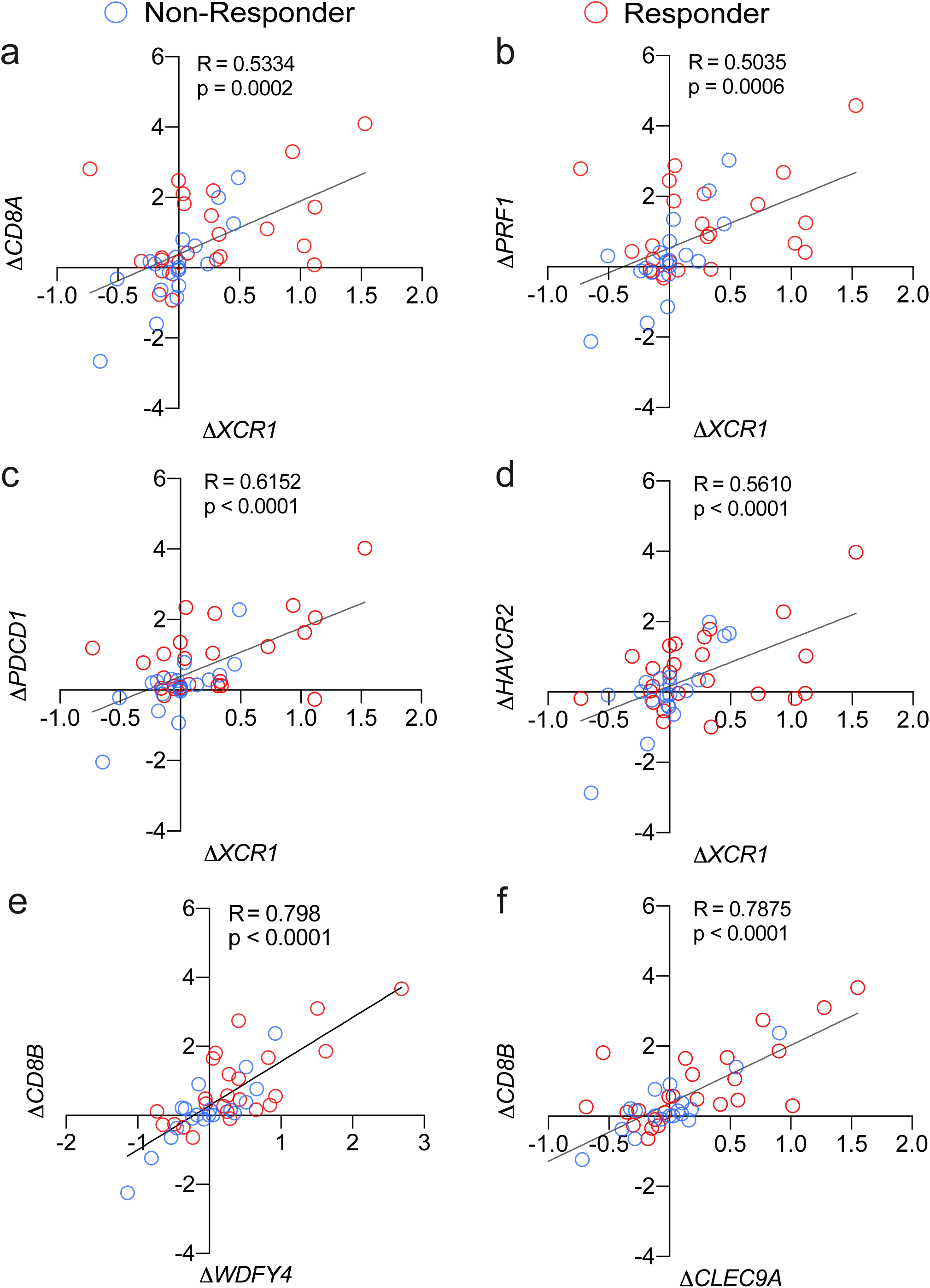
Correlation between fold changes in the level of cDC1 markers and that of CD8 T cell markers before and after αPD-1 treatment in melanoma patients. Fold change (Δ) of gene expression was calculated as log2(FPKM_Post-treatment_ plus pseudo-count of 0.1) minus log2(FPKM_Pre-treatment_ plus pseudo-count of 0.1). Pair-wise Pearson’s correlation between Δ*XCR1* and Δ*CD8A* (**a**), Δ*XCR1* and Δ*PRF1* (**b**), Δ*XCR1* and Δ*PDCD1* (**c**), Δ*XCR1* and Δ*HAVCR2* (**d**), Δ*WDFY4* and Δ*CD8B* (**e**), Δ*CLEC9A* and *ΔCD8B* (**f**) were calculated. Correlation coefficients were also indicated.

**Supplementary Figure 9.**
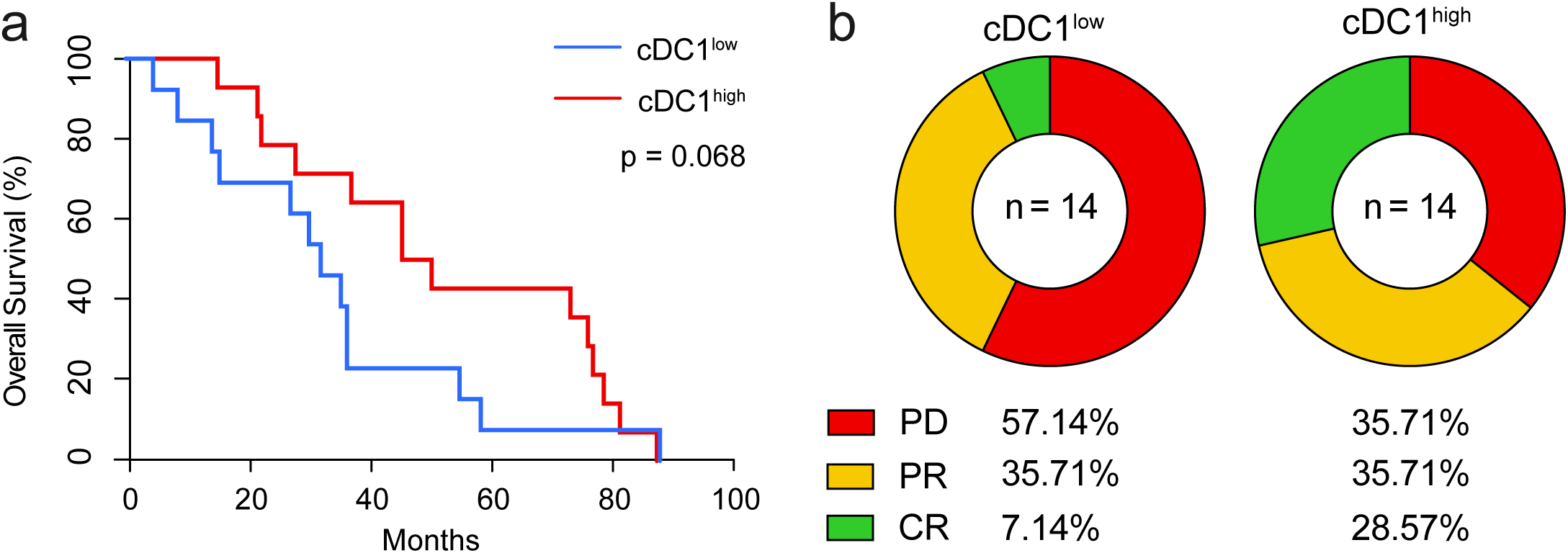
Association between pre-treatment cDC1 signature expression and survival of metastatic melanoma patients receiving αPD-1 therapy. **a**, Association between pre-treatment tumor cDC1 signature expression and survival in metastatic melanoma patients treated with αPD-1 antibody therapy. cDC1 signature is calculated as the average expression of *XCR1*, *WDFY4*, *IRF8*, *TLR3*, and *CLEC9A*. Survival curves were compared using Gehan–Breslow–Wilcoxon test which assigns extra weight for early time points. **b**, Distribution of clinical responses to αPD-1 therapy stratified by expression level of pre-treatment cDC1 signature. PD, progressive disease; PR, partial response; CR, complete response.

